# Live imaging in zebrafish reveals tissue-specific strategies for amoeboid migration

**DOI:** 10.1101/2024.08.14.607647

**Authors:** Tanner F. Robertson, Jon Schrope, Zoe Zwick, Julie K. Rindy, Adam Horn, Anna Huttenlocher

## Abstract

Amoeboid cells like leukocytes can enter and migrate within virtually every tissue of the body, even though tissues vary widely in their chemical and mechanical composition. Here, we imaged motile T cells as they colonized peripheral tissues during zebrafish development to ask if cells tailor their migration strategy to their local tissue environment. We found that T cells in most sites migrated with f-actin-rich leading-edge pseudopods, matching how they migrate *in vitro*. T cells notably deviated from this strategy in the epidermis, where they instead migrated using a rearward concentration of f-actin and stable leading-edge blebs. This mode of migration occurs under planar confinement *in vitro*, and we correspondingly found the stratified keratinocyte layers of the epidermis impose planar-like confinement on leukocytes *in vivo*. By imaging the same cell type across the body, our data collectively indicates that cells adapt their migration strategy to navigate different tissue geometries *in vivo*.

## Introduction

Cell migration plays a central role in immune function, development, and cancer invasion (1–3). A growing body of *in vitro* studies have found that cells sense the chemical and mechanical composition of their environment and modify their migration strategy to effectively navigate these spaces (1, 4, 5). In some contexts, cells employ leading edge actin polymerization in the form of lamellipodia or pseudopods to effectively migrate (6, 7). In other contexts, cells may harness intracellular pressure to move via leading edge blebbing (8–10). Migratory plasticity is thought to be critical because tissues vary widely in their composition of cells, extracellular matrix (ECM) components, adhesive ligands, chemoattractants, and other migration-influencing factors (1, 4, 11). Movement across the body by leukocytes or disseminating cancer cells may therefore depend on toggling between different migration modes as they encounter different tissue environments. However, direct comparisons of cell migration strategies across distinct tissue environments *in vivo* are lacking.

During vertebrate development, leukocyte populations enter and colonize a wide variety of tissues (12–14), providing an opportunity to monitor cell migration in heterogenous tissues. Here, we took advantage of the transparent nature of larval zebrafish to directly compare amoeboid leukocyte migration within multiple tissue environments *in vivo* (15). We found that T cells spontaneously migrated in the thymus, intestines, and epidermis during larval development and generated an f-actin reporter line to compare migration strategies in each site. In line with previous studies investigating T cell f-actin dynamics within collagen matrices (16) and murine lymph nodes (17), we found that T cells in the thymus and intestines migrated with an accumulation of leading edge f-actin. In contrast, T cells in the epidermis migrated with f-actin concentrated in the rear of the cell and nearly devoid from leading edge. We found that these epidermal T cells used retrograde actin flow and myosin-dependent contractility to move, mirroring the “stable bleb” mode of migration (also called leader bleb and A2 migration) employed by a variety of highly contractile cells under planar confinement (18–21). In accordance with these studies, we provide evidence that epidermal T cells are highly confined between keratinocyte layers as they migrate *in vivo*. These data support the idea that leukocytes use tissue-specific migration programs to effectively navigate the host.

## Results

### Characterization of an epidermal T cell population in adult zebrafish

We recently identified a body-spanning lymphoid tissue in zebrafish that localizes to the pocket formed by overlapping scales (22). Using pan-T cell reporter *lck*:GFP adult zebrafish, we found that zebrafish also maintain a population of epidermal T cells on the outer scale layer (Fig. 1A). To determine if scale and TLN T cells represent distinct populations, we performed high resolution imaging of both groups. We found that scale T cells maintain a dendritic cell shape, which distinguished them from TLN T cells, which maintain a compact rounded morphology more typical of lymphocytes (Fig. 1B-C). This difference in morphology was confirmed by measuring solidity (cell area/convex hull area), which is lower in cells with irregular, protrusive shapes (Fig. 1D). In line with our previous report, we found that *cd4.1*, which labels helper T cells (23), was expressed by the majority of TLN T cells but relatively few scale T cells (Fig. 1B, C, E). Scale T cells did, however, form a network with Langerhans cells (LCs), an epidermis-resident macrophage population that highly express *cd4.1* in zebrafish (Fig. 1B) (23–25). Further, we found that scale T cells were sessile under homeostatic conditions, which contrasts them with the constitutively motile TLN T cells (Fig. 1G). When adult zebrafish are descaled, TLN T cells are released into suspension (22), but we found that the scale T cell-LC network remains scale-bound (Fig. 1H), in line with previous reports documenting T cells on plucked scales (24–27). These data indicate that scale T cells and TLN T cells are distinct populations.

**Figure 1.**
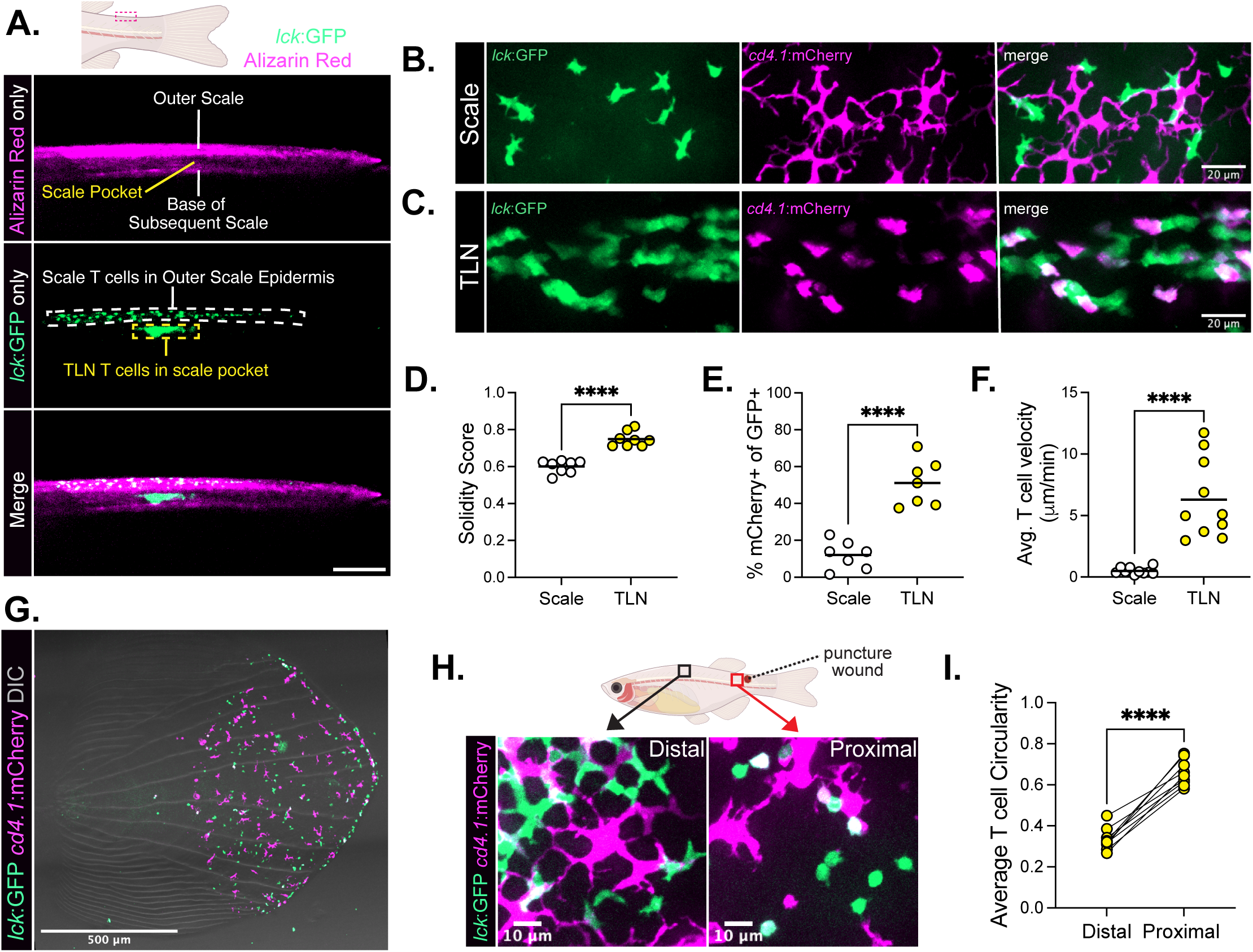
Characterization of a dendritic epidermal T cell population in adult zebrafish. **A.** Imaging of a tg(lck:GFP) adult zebrafish stained with alizarin red shows TLN T cells in the scale pocket and a separate population of T cells on the outer scale. Scale bar = 100 microns. **B-C.** Representative images of outer scale (B) and TLN (C) regions in a tg(lck:GFP; cd4.1:mCherry) adult zebrafish. Scale bar = 20 microns. **D-F.** Quantification of the solidity score (cell area divided by convex hull area; D), percent cd4.1 co-expression among GFP+ T cells (E), and average T cell velocity (F) for scale and TLN T cells, with each dot corresponding to a fish pooled from three independent experiments. Unpaired t-test, ****p<0.0001. **G.** Imaging of a scale plucked from an adult tg(lck:GFP; cd4.1:mCherry) zebrafish showing outer scale T cells and Langerhans cells remaining scale-bound. Scale bar = 500 microns. **H-I.** Representative images (H) and associated quantification (I) of the circularity (4pi(area/perimeter^2)) of T cells distal and proximal to a puncture wound administered 2 hours prior, with lines connecting measurements from the same fish. Scale bar = 10 microns. Paired t-test, ****p<0.0001.

In our initial characterization of scale T cells, we noticed they shared many features with a specialized T cell population in mice called dendritic epidermal T cells (DETCs). Murine DETCs also maintain a dendritic morphology, lack CD4 co-expression, and are sessile under homeostatic conditions (28–33). One defining characteristic of murine DETCs is that they respond to nearby tissue damage by retracting their protrusions and rounding up (32). To determine if zebrafish scale T cells behaved similarly, we induced puncture wounds in adult tg(*lck*:GFP; *cd4.1*:mCherry) zebrafish and imaged proximal and distal T cell morphology 2 hours later (Fig. 1I). We found that, like murine DETCs, zebrafish scale T cells also rounded up in response to nearby tissue damage (Fig. 1I-J). Collectively, these data indicate that zebrafish scale T cells may be analogs of murine DETCs.

### T cells form an epidermal network with Langerhans Cells during larval development

During embryonic development in mice, DETC precursors are the first T cells to egress from the thymus and subsequently traffic to the developing epidermis (12, 13). To determine the degree of similarity between zebrafish scale T cells and murine DETCs, we investigated when scale precursor T cells first emerged during zebrafish development. In line with previous studies (34, 35), we found that the thymus first appears between 3-5 dpf near the developing gills, which are visible in the endothelial *kdrl* blood vessel reporter line (Fig. 2A)(36). At 5 dpf, T cells began accumulating along the caudal vein of the tail fin (Fig. 2A). In 2-hour live imaging experiments with acquisition rates rapid enough to visualize circulating T cells in 6 dpf larvae, we found that T cells are captured from circulation onto the caudal vein at a rate of 1.20 ± 1.81 cells per hour (Fig. S1A-C). Some T cells dislodged from the caudal vein and returned to circulation resulting in a lower transmigration rate of 0.70 ± 0.67 cells per hour (Fig. S1A, D). T cell capture and transmigration occurred preferentially at the caudal vein over the other blood vessels present in the caudal peduncle (Fig. S1A-D). T cells entering the tail fin are therefore among the earliest thymic emigrants in zebrafish larvae.

**Figure 2.**
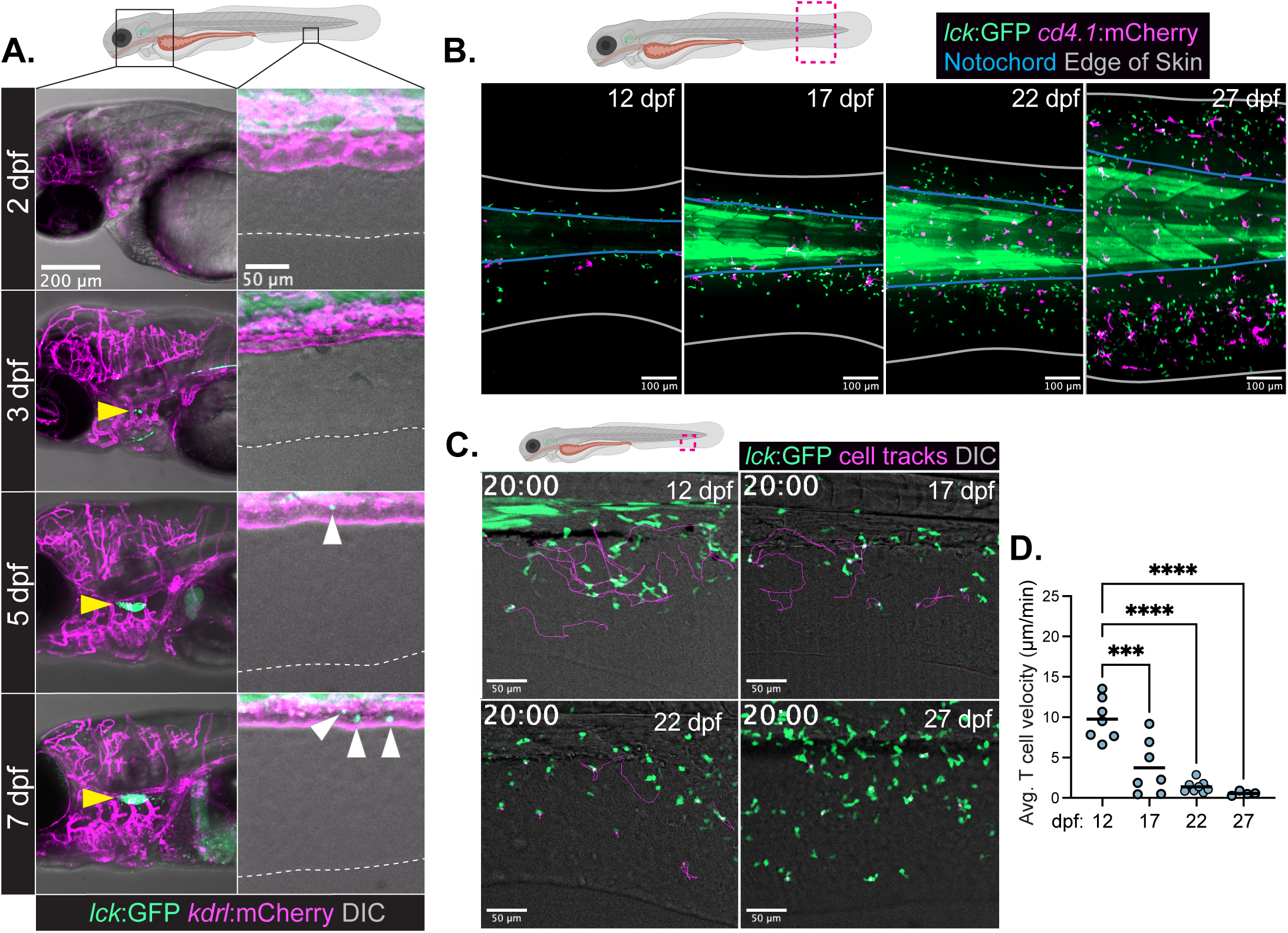
The T cell-Langerhans Cell Network forms during larval development. **A.** Imaging of a representative tg(lck:GFP; kdrl:mCherry) larval zebrafish head and indicated skin region at 2, 3, 5, and 7 dpf showing the development of the thymus (yellow arrowheads) and individual T cells accumulating along the caudal vein (white arrowheads) with the white dotted line marking the edge of the skin. Representative of 9-12 larvae per timepoint from 3 independent clutches and experiments. Head scale bar = 200 microns, skin scale bar = 50 microns. **B.** Imaging of the caudal peduncle of tg(lck:GFP; cd4.1:mCherry) larval zebrafish between 12 and 27 dpf showing T cells (green) and Langerhans cells (magenta) spreading across the skin. Blue outline marks the notochord, white outline marks the edge of the skin. Representative of 6-8 fish from at least 3 independent clutches and experiments. Scale bar = 100 microns. **C-D.** Spontaneous T cell (green) migration at 12, 17, 22, and 27 dpf shown with overlaid cell tracks (magenta) (C) and corresponding quantification (D) of average T cell velocity of individual fish (blue dots) with the mean at 12 dpf compared to every other mean with an ordinary one-way ANOVA. ***p=0.0002, ****p<0.0001. Scale bar = 50 microns.

By 12 dpf, T cells have begun accumulating in the skin of the tail fin, but remain near the notochord, and caudal vein (Fig. 2B). Over the course of larval development, which ends at approximately 30 dpf, these T cells progressively spread out across the skin, covering it in a network of T cells (Fig. 2B). At 12 dpf, these T cells are highly motile, but they become progressively sessile as the T cell network forms (Fig. 2C-D, Videos S1). LCs spread out across the skin over a similar developmental timeframe (Fig. 2B). This sessile T cell-LC network is therefore established by the end of larval development and precedes the formation of either the TLN or scales themselves, both of which emerge during the juvenile stages of development (22, 37). Collectively, these data indicate that scale T cell precursors are among the first thymic emigrants in zebrafish, mirroring DETCs in mice.

### Epidermal T cells migrate with f-actin-poor leading edges during development

Motile cells can adopt different strategies to migrate and will often tailor their migratory response to their environment (1, 4, 5). Being able to toggle between different modes of migration likely enables leukocyte and cancer cell migration across distinct and heterogenous tissues. However direct comparisons of cell migration strategies across diverse tissue environments *in vivo* are lacking. In addition to DETCs, additional T cell populations leave the thymus and colonize peripheral tissues during embryonic and perinatal development in mice (12). Because these processes occur largely *in utero* in mice, visualizing T cell migration across these tissues is technically challenging. We therefore wondered if T cells also colonized additional tissues in zebrafish and whether we could use this model organism to compare amoeboid migration strategies in distinct tissues *in vivo*.

In whole-fish confocal tiles at 12 dpf, T cells could be easily identified in the thymus (Fig. 3A), where their migration has been previously investigated (23). We also consistently observed T cells in the intestines (Fig. 3A), which could be distinguished from surrounding tissue by peristaltic movement. To first ask if T cells engaged in tissue-specific migration strategies, we generated a T cell specific f-actin reporter by injecting 1-cell stage *lck*:GFP embryos with a construct containing lifeact-ruby (38) downstream of the T cell specific *lck* promoter (Fig. 3B) (34). We screened for fish with incomplete lifeact-ruby expression in the thymus, as this made tracking cells in the thymus feasible (Fig. 3C). We performed live imaging of T cell migration in each site and compared f-actin dynamics and organization (Fig. 3D). We found that the total lifeact-ruby signal was significantly higher in the thymus compared to peripheral tissues, which required that we contrast and scale images on a cell-by-cell basis to compare their f-actin organization (Fig. S2A-B). We found that cells in the thymus and intestines migrated with f-actin-rich pseudopods at the leading edge (Fig. 3D, Video S2), mirroring how amoeboid cells migrate in extracellular matrices *in vitro* and a wide variety of non-epithelial tissues *in vivo* (16, 17, 39–41). In contrast, skin T cells migrated with their f-actin concentrated in the rear of the cell and nearly devoid from the leading edge (Fig. 3D, Video S2). We quantified f-actin organization by calculating the ratio of leading edge (LE) to trailing edge (TE) f-actin and found T cells in the skin had significantly less leading-edge f-actin than their counterparts in the thymus and intestines (Fig. 3E). Skin T cells also migrated with a highly elongated morphology relative to motile T cells in the thymus and intestines (Fig. 3D, F). We performed line profile analysis of T cells at the low, middle, and high end of the LE/TE f-actin ratio spectrum (Fig. 3G-I). At the low end (LE/TE = 0.16), cells had a cytoplasmic bulge at the leading edge largely devoid of f-actin (Fig. 3G). At the middle (LE/TE = 0.36) and high (LE/TE=0.79) ends of the spectrum, T cells maintained relatively small concentrations of f-actin at the leading edge, but the GFP+ cytoplasm extended past the f-actin, suggesting f-actin polymerization was not driving leading edge expansion (Fig. 3H-I). Within the thymus, the high cellular density precluded this kind of analysis, but for intestinal T cells we did not observe cytoplasm bulging past the leading-edge (Fig. 3J). These data indicate that skin T cells are utilizing a different migration strategy than T cells in the thymus and intestines.

**Figure 3.**
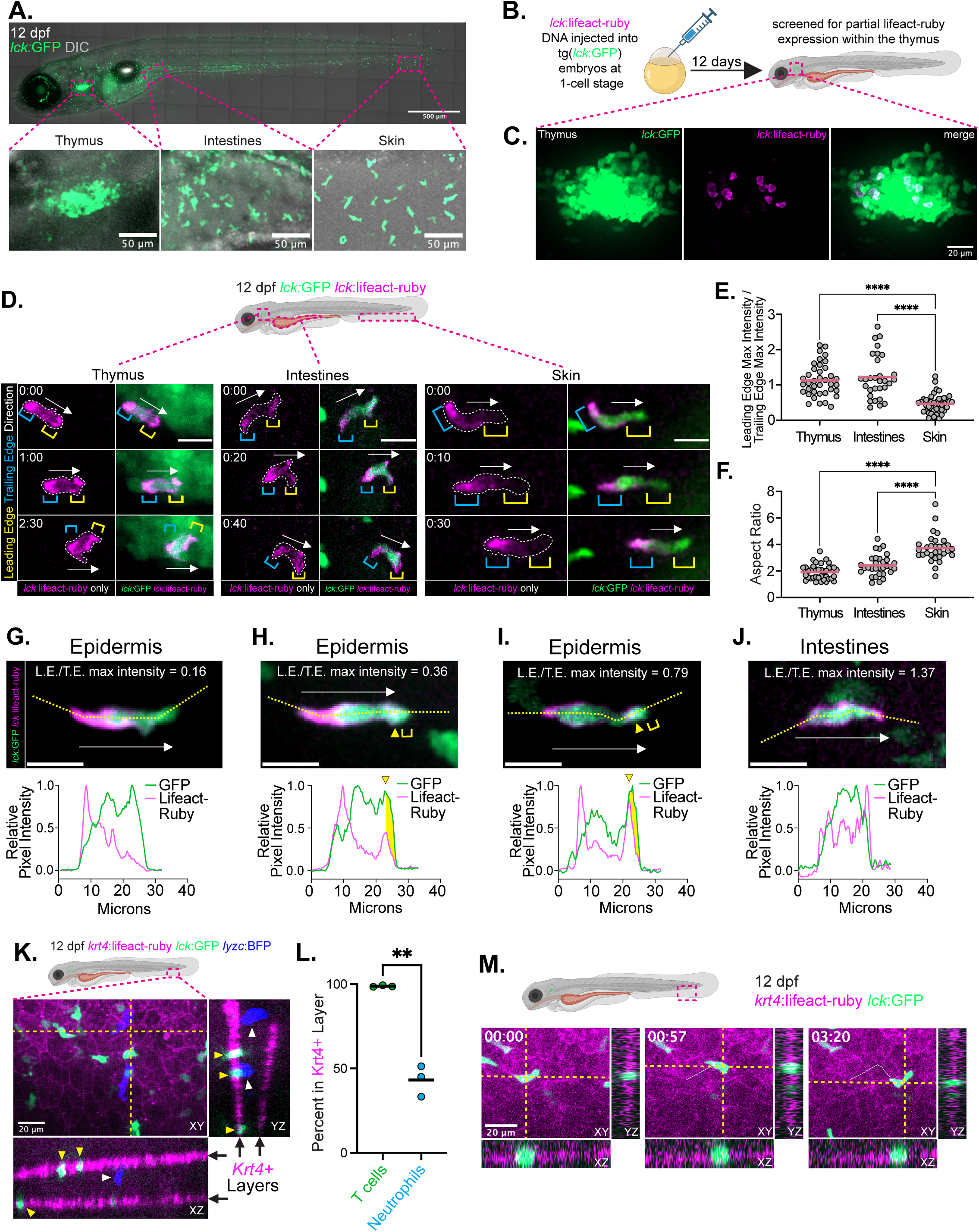
T cell migrate with f-actin-poor leading edges in the larval epidermis. **A.** The distribution of T cells across the body in a 12 dpf tg(lck:GFP) larvae, with insets highlighting T cells in the thymus, intestines, and epidermis. Representative of 4 whole-fish tiles from 3 independent experiments and clutches. **B.** Experimental design for the generation of F0 12 dpf zebrafish larvae with lifeact-expressing T cells. **C.** A confocal z-stack of an F0 tg(lck:GFP; lck:lifeact-ruby) 12 dpf thymus showing the partial expression of lifeact among developing T cells. Representative of 10 thymi pooled from 3 independent experiments. **D.** Representative images of GFP+ and lifeact-ruby+ T cells in the thymus (42 cells from 8 larvae), intestines (30 cells from 10 larvae), and epidermis (42 cells from 11 larvae) from 11-12 dpf larvae pooled from 3 independent experiments with arrows indicating the instantaneous direction of migration and with the leading edge and trailing edge noted. The isolated lifeact-ruby channel is provided with the cell outlined to highlight the distribution of f-actin. **E-F.** Quantification of the ratio of lifeact-ruby max intensity in the leading edge (front third of cell) and trailing edge (rear third of cell) (E) and aspect ratio (F) with each dot representing a cell pooled from three independent experiment. The skin mean is compared with the other means with an ordinary one-way ANOVA. ****p<0.000. **G-I.** Images of epidermal T cells from tg(lck:GFP; lck:lifeact-ruby) and corresponding line profile analysis showing either the absence of f-actin at the leading edge (G) or GFP+ cytoplasm extending past f-actin (yellow region; H and I). **J.** Images of an intestinal T cell and accompanying line profile analysis as in C-E showing f-actin at the leading edge. **K.** T cells occupy the epidermis in larval zebrafish. A z-stack and corresponding orthogonal views of a 12 dpf tg(krt4:lifeact-ruby; lck:GFP; lyzc:BFP) showing the krt4+ epidermis (magenta), T cells (green), and neutrophils (blue). Scale bar = 20 m **L.** Quantification of the percentage of skin T cells and neutrophils within the krt4+ epidermal layers. Each dot represents the average of 3-6 larvae from separate clutches and experiments. Means were compared with a paired t-test. **p=0.0088. **M.** Timelapse imaging and the corresponding orthogonal projections of a T cell (green) migrating in the epidermis (magenta) of a 11-12 dpf tg(krt4:lifeact-ruby; lck:GFP) larvae. Representative of 52 T cells from 7 larvae and three independent experiments.

To understand why T cells were migrating differently in the skin than the other tissues, we next investigated the tissue environment that skin T cells were migrating in. Previous studies have visualized amoeboid migration within the tail fin skin of larval zebrafish (7, 42–45), but whether these cells migrate in the extracellular matrix-rich dermis, cellularly dense epidermis, or a different layer of the tissue remains poorly understood. The epidermis of larval zebrafish is a bi-layered stratified epithelial tissue comprised of basal and suprabasal keratinocytes, which can be visualized with transgenic krt4 reporter fish (46). We generated triple reporter tg(*krt4*:lifeact-ruby; *lck*:GFP; *lyzC*:BFP) fish to simultaneously label the epidermis (magenta), T cells (green), and neutrophils (blue; Fig. 3K). In *z*-stacks of the larval skin, T cells were primarily found in the krt4+ epidermis (Fig. 3K-L). This was not because other layers of the skin were inaccessible to leukocytes, as neutrophils occupied both the krt4+ epidermis and deeper dermal layers in the same fish (Fig. 3K-L). To determine if T cells were moving between the krt4+ epidermis and other skin regions as they migrated, we visualized T cell migration in orthogonal projections and found that T cells remained localized to the epidermis as they moved (Fig. 3M).

### Epidermal T cells migrate with f-actin-poor leading edges in scale explants

We next investigated if f-actin-poor leading-edge migration by T cells was restricted to early larval development or if it instead represented a common strategy for navigating the epidermis in larvae and adults alike. In adult zebrafish, the epidermis wraps around the scale and remains attached following plucking, which allows for *in situ* monitoring of epidermal leukocyte behavior (24, 47). We found that when scales were carefully isolated to minimize epidermal damage (Fig. S3), T cells were mostly sessile in scale explants (Fig. 4A-B, Video S3), mirroring their behavior in scales attached to the host (Fig. 1G). However, when we used forceps to damage the epidermis (Fig. S3), T cells became significantly more motile (Fig. 4A-B, Video S3). Lifeact-expressing T cells exhibited a wide range of velocities in wounded scale explants, from sessile to nearly 30 μm/min (Fig. 4C-E). In rapidly migrating cells, we observed a clear rearward concentration of f-actin (Fig. 4C, Video S4), mirroring how T cells migrate in the larval epidermis (Fig. 3D, Video S2). In slowly migrating cells, this strong rearward f-actin concentration was lost (Fig. 4D). We observed a strong correlation between the absence of f-actin at the leading edge and velocity (Fig. 4E). That is, the fastest epidermal T cells maintained the lowest LE/TE f-actin ratio resulting in a strong negative correlation (R^2^ = 0.6204) between these two parameters (Fig. 4E, Video S4). In some cells, f-actin would transiently accumulate at the leading edge, but this was associated with cells stopping and changing direction (Fig. 4F). Collectively, these data indicate that f-actin-poor leading edges are a common feature of rapidly migrating epidermal T cells irrespective of developmental stage.

**Figure 4.**
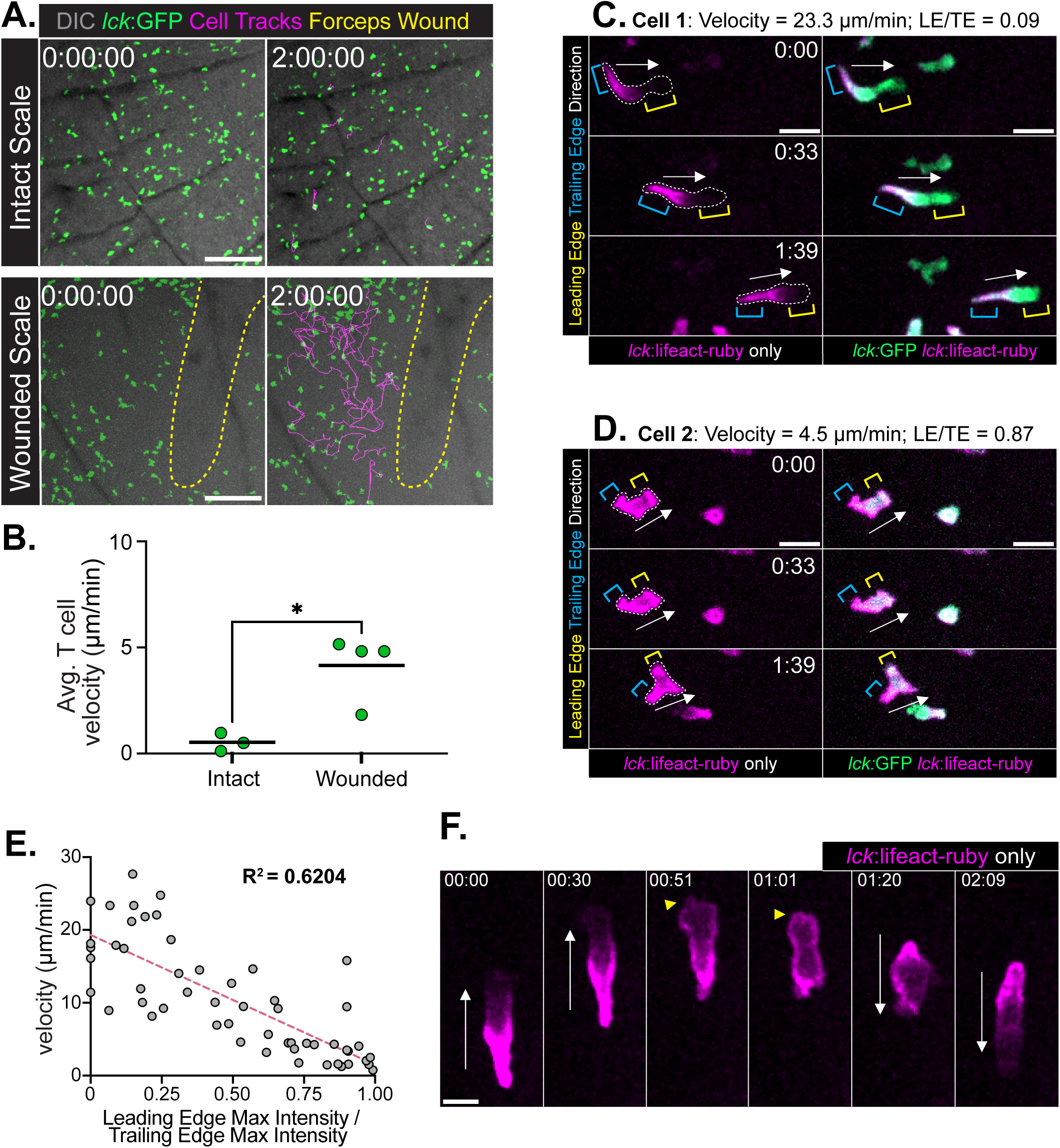
T cells migrate with f-actin-poor leading edges in scale explants. **A-B.** Timelapse imaging of an intact and forceps-wounded (yellow dotted line) scale with overlaid cell tracks (magenta) (A) and associated quantification of T cell velocity (B), with each dot corresponding to the average T cell velocity from a scale explant from 3-4 independent experiments. Unpaired t-test, *p=0.0122. Scale bar = 100 microns. **C-D.** Timelapse images of a fast and slow GFP+ and lifeact-ruby+ T cell in scale explants with the LE:TE max lifeact intensity ratio and velocity noted. Scale bar = 10 microns. **E.** T cell velocity plotted against the LE:TE lifeact max intensity ratio showing the negative correlation between speed and leading-edge f-actin for scale explant T cells. Each point represents a single cell (5 cells tracked in 3 independent movies over 3 independent experiments; 45 cells total). **F.** Timelapse imaging of a scale explant from a tg(lck:lifeact-ruby) fish showing leading-edge f-actin accumulation (yellow arrowheads) corresponding with the T cell stopping and changing directions (noted with white arrows). Scale bar = 5 microns.

### Epidermal T cells migrate with stable leading-edge blebs

Leukocytes, including T cells, can migrate through leading edge blebbing where actomyosin contractility drives cytoplasmic protrusions past the actin cortex (8). Leading edge blebs can either be intermittent or stable (10). Intermittent blebs form rapidly when intracellular pressure causes the cytoplasm to extend past the actin cortex (48). When the cortex reforms, a subsequent bleb can be generated, allowing the cell to migrate through continuous intermittent blebbing (10, 48–50). In the stable bleb mode of migration, cells form a single large cytoplasmic bleb at the leading edge and propel themselves through retrograde actin flow and myosin-dependent contraction (18–21). Intermittent blebs form rapidly, so we imaged tg(*lck*:GFP; *lck*:lifeact-ruby) T cells with ∼1.2 second acquisition rates in the epidermis (Fig. S4A). We did not observe rapid blebbing at the leading edge; the cytoplasmic protrusions appeared highly stable (Fig. S4A). We did, in rare instances, observe intermittent blebs in epidermal T cells, but these typically occurred outside the leading edge and collapsed back into the cell body without notably impacting cell migration (Fig. S4B). These data suggest that epidermal T cells are migrating via stable rather than intermittent blebs.

### Epidermal T cells utilize retrograde actin flow and contractility to rapidly migrate

The rearward concentration of f-actin and leading-edge bulge of cytoplasm observed in motile epidermal T cells are consistent with the stable bleb mode of migration (18–21). This migration strategy occurs under planar confinement *in vitro*, when cells are sandwiched between adjacent surfaces (also called 2.5D confinement) and relies upon retrograde actin flow and myosin-dependent contractility to propel the cell forward (18–21, 51). We therefore asked whether epidermal T cells were using retrograde actin flow and myosin-dependent contractility for rapid migration *in vivo*. Retrograde actin flow is typically visualized by total internal reflection fluorescence (TIRF) microscopy, which preferentially illuminates fluorophores within a few hundred nanometers from the sample-coverslip interface (52, 53). Scale explants provide an *in situ* imaging opportunity, but the T cells are too far from the imaging surface to use TIRF microscopy. As an alternative, we performed rapid acquisition, high resolution confocal microscopy on lifeact-expressing epidermal T cells utilizing the stable bleb migration strategy *in situ* (Fig. 5A). In some cells, the f-actin signal was too homogenous to analyze, but we were able to analyze actin dynamics in 15 cells pooled from three independent experiments (Fig. 5A-D). These images revealed that f-actin at the leading edge to midbody of the cell stays in the same absolute position and moved retrograde relative to the leading edge of the cell (Fig. 5B-D), matching the behavior of T cells that migrate *in vitro* under 2.5D confinement (51).

**Figure 5.**
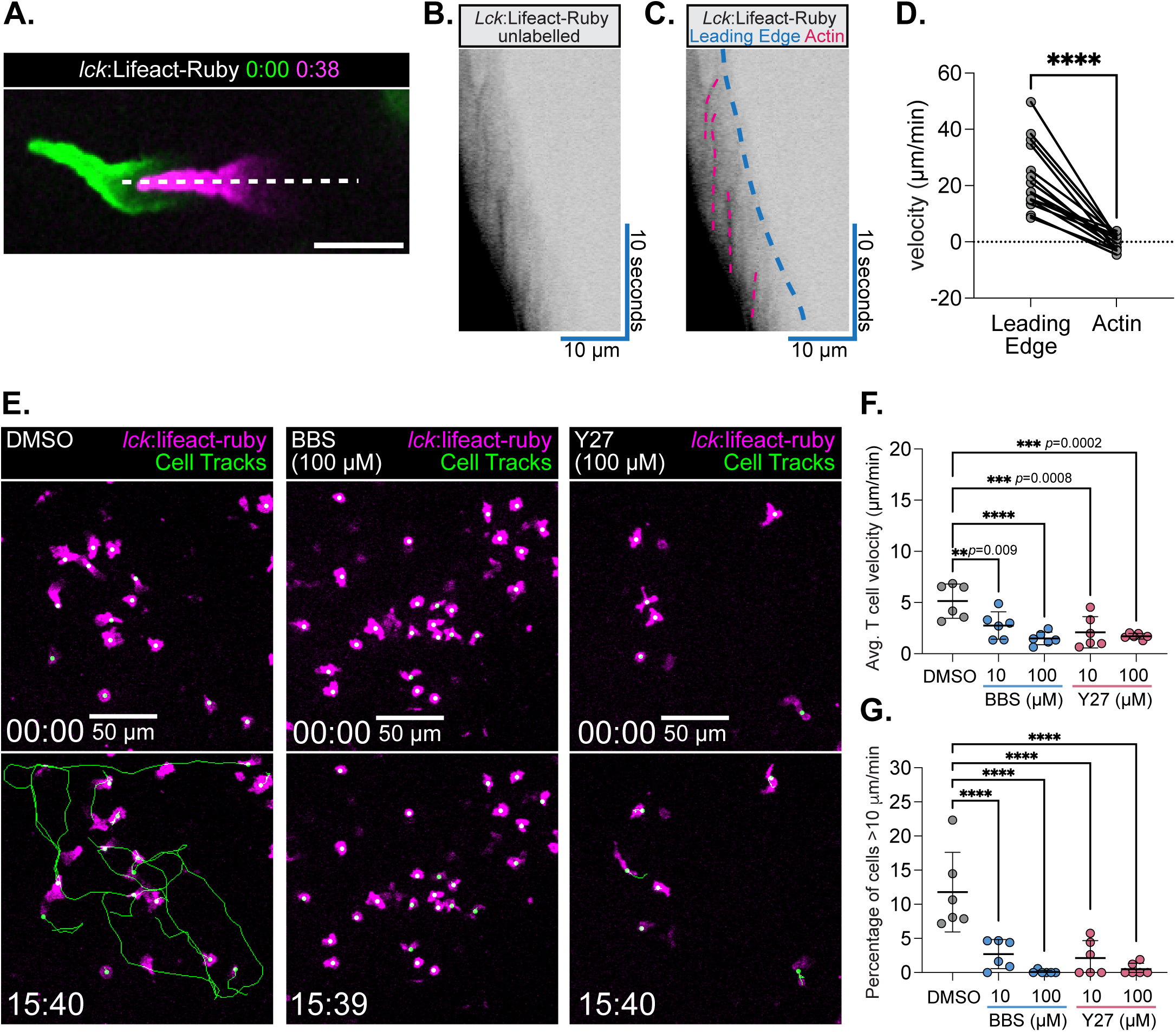
T cells use retrograde actin flow and myosin-dependent contractility to rapidly migrate through the epidermis. **A.** Snapshots of a motile lifeact-expressing T cell at the indicated timepoints captured by confocal microscopy. **B-C.** A kymograph generated from the line in (A) left unlabelled (B) or with the leading edge and actin traces labelled (C). **D.** Mean leading edge and actin velocities for 15 cells, represented by dots, pooled from three independent experiments. Means compared with a paired t-test; ****p<0.0001. **E-G.** Timelapse imaging and overlaid cell tracks (green) of lifeact+ T cells (magenta) migrating in scale explants treated with DMSO or the indicated concentration of BBS or Y27 shown with the corresponding quantification of average T cell velocity (F) and the percentage of T cells whose average velocity exceeds 10 m/s (G). Each point represents the mean of a single scale explant replicate, data pooled equally over three independent experiments. Time in mm:ss, Scale bar = 50 m. One-way ANOVA comparing the DMSO mean to all other means. ****p<0.0001, other p-values noted on graph.

We next investigated whether actomyosin contractility was required for this rapid epidermal migration. We incubated scale explants with the myosin inhibitor blebbistatin (BBS) or the Rho-associated Protein Kinase (ROCK) inhibitor Y-27632 (Y27) and measured the impact of these drugs on epidermal T cell migration (54, 55). Both BBS and Y27 treatment resulted in a significant decrease in the average T cell velocity in wounded scale explants (Fig. 5E-F, Video S5). The percentage of cells migrating rapidly (>10 μm/min) dropped in a dose-dependent manner following BBS and Y27 treatment (Fig. 5G). The rearward f-actin concentration, retrograde actin flow, leading-edge cytoplasmic bulge, and dependence on actomyosin contractility collectively indicate that epidermal zebrafish T cells are utilizing stable bleb migration to rapidly move within the epidermis.

### The epidermis imposes 2D planar confinement on epidermal T cells

A wide variety of cells, including T cells and other leukocytes, employ stable bleb migration under planar confinement *in vitro* (18–21, 51). Given that T cells use stable blebs in the epidermis but not the thymus or intestines, we asked whether the epidermis imposes planar confinement on T cells *in vivo*. We initially hypothesized that T cells may be migrating between adjacent keratinocytes within either the basal or suprabasal keratinocyte layers, but T cells did not appear to follow the intercellular paths visible in *krt4*:lifeact-ruby reporter fish (Fig. 6A). We next hypothesized that T cells may be sandwiched between the basal and suprabasal keratinocyte layers. In *krt4* reporter fish, we found it challenging to resolve the basal and suprabasal layers in orthogonal projections when fish were mounted on their side (Fig. 3K). We instead mounted and visualized larval zebrafish upright, such that we could simultaneously visualize epidermal T cells and distinguish the basal and suprabasal keratinocyte layers in the XY rather than Z planes (Fig. 4B). In live imaging experiments and corresponding line profile analysis on upright mounted larval zebrafish, we found that T cells migrated in apparent confinement between the basal and suprabasal keratinocyte layers (Fig. 6B-C). We found that epidermal T cells had a maximum *z*-height of ∼3 microns as they migrated *in vivo* (Fig. 6D), matching the degree of confinement needed to drive stable bleb migration *in vitro* (18). While there is no single or direct measurement of confinement, cells under planar confinement increase their 2D surface area as they flatten (56, 57). We therefore measured the 2D surface area in motile T cells in the thymus, intestines, and epidermis of flat-mounted fish as a proxy for confinement and found that epidermal T cells had a significantly higher 2D surface area (Fig. 3D, Fig. 6E). These data are consistent with the epidermis imposing planar confinement on T cells *in vivo* to elicit the same kind of stable bleb migration caused by planar confinement *in vitro*.

**Figure 6.**
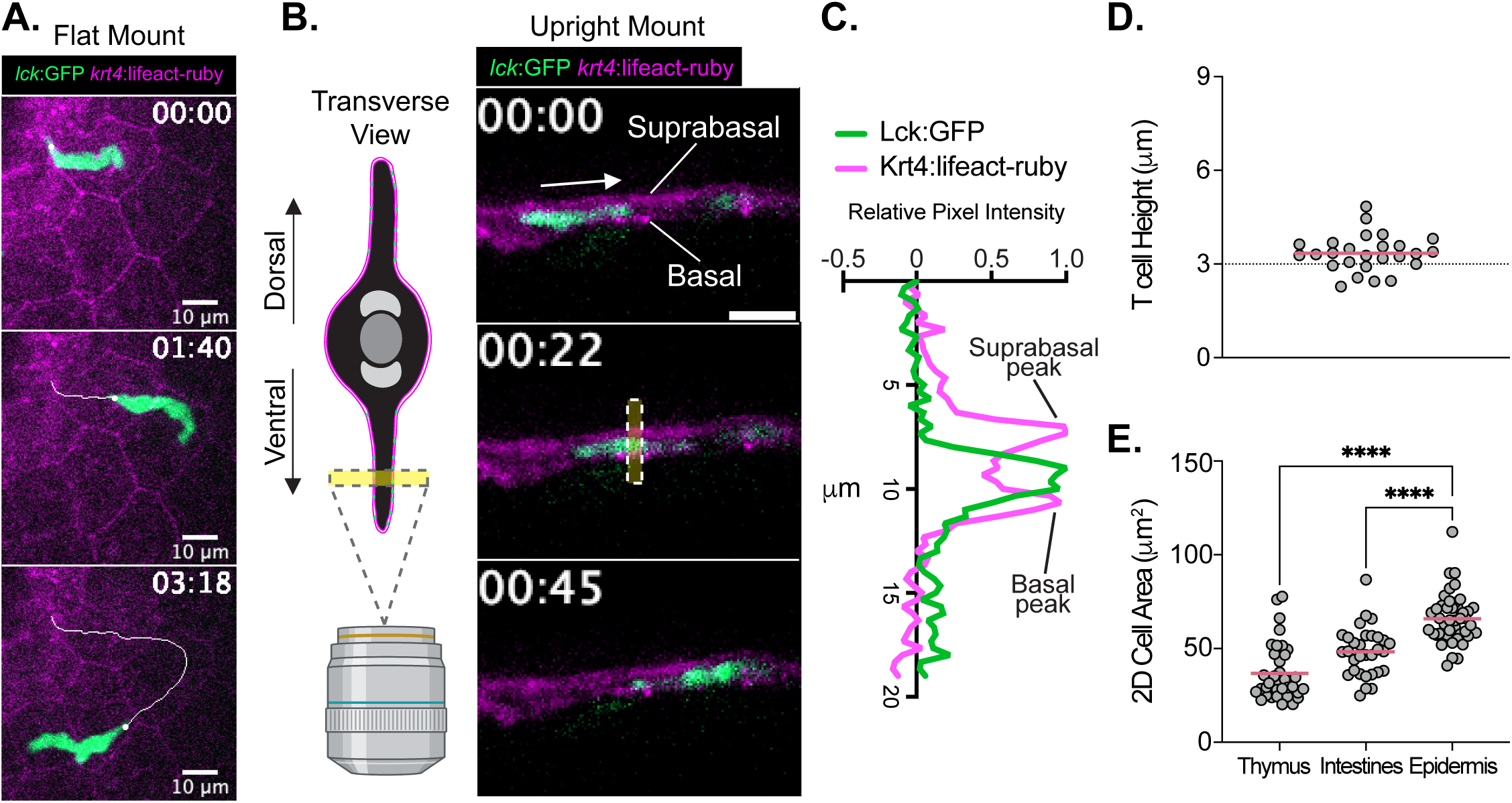
The epidermis imposes planar confinement on motile T cells. **A.** Single z-plane timelapse imaging of tg(lck:GFP; krt4:lifeact-ruby) zebrafish showing a T cell migration path not occurring between adjacent keratinocytes. Representative of 3 independent experiments. Scale bar = 10 microns. **B.** A schematic for how larval zebrafish are mounted and imaged upright and a timelapse image showing a T cell migrating between the basal and suprabasal keratinocyte layers. Representative of 3 independent experiments. Scale bar = 10 microns. **C.** A line profile of the highlighted region in the middle frame of (B.) showing the T cell fluorescent peak between the peaks corresponding to the basal and suprabasal keratinocyte layers. **D.** The maximum height of T cells migrating between the basal and suprabasal keratinocyte layers measured in upright mounted larvae. Each dot represents a cell pooled from 3 independent experiments. **E.** The 2D area of T cells in the corresponding tissue measured in flat-mounted fish, with each dot corresponding to a cell pooled from 3 independent experiments. Means are compared with an ordinary one-way ANOVA. **p=0.0025, ****p<0.0001.

## Discussion

In this study we used zebrafish to study cell migration strategies in different tissue environments *in situ* and found that T cells migrate via stable blebs in the epidermis but not the thymus or intestines. This supports a growing body of *in vitro* studies demonstrating that amoeboid cells sense and tailor their migration strategy to their environment to efficiently move through distinct interstitial spaces. The emerging picture from these studies is that cells migrating with high contractility in confining, low-adhesive environments utilize stable bleb migration, and investigators have identified numerous cytoskeletal regulators critical for this migration (18–21, 58–61). However, the *in vivo* relevance of such a migration strategy remains unclear. To our knowledge, the only examples of cells utilizing stable bleb migration *in vivo* are zebrafish progenitor cells with artificially increased contractility (19) and a melanoma sub-line with enhanced myosin activity (62). Epidermal zebrafish T cells may therefore represent the first example of cells utilizing stable bleb migration spontaneously *in vivo* and may offer a useful system for interrogating this mode of migration under physiological conditions. By dissecting the different molecular requirements for migration through different tissues, investigators may eventually be able to enhance or inhibit leukocyte trafficking to and movement within specific tissues for therapeutic purposes.

As *in vitro* approaches to study cell migration become increasingly elegant, investigators should consider how their devices and environment reflect *in vivo* tissue geometries. *In vitro* approaches typically employ one of three kinds of environments: 2D surfaces coated with an adhesive ligand, 2D planar confinement, and 3D collagen-based matrices (63, 64). Early investigation into cell migration often employed 2D surfaces, which overemphasize the role of specific adhesive receptors like integrins. This is why finding that leukocytes can migrate *in vivo* in the absence of integrins was such a paradigm-shifting finding in the field (40). Most tissues of the body have a significant connective tissue component composed of fibrillar collagen and other extracellular matrix (ECM) elements, and studies using *in vitro* collagen matrices likely approximate this kind of tissue composition (11, 65). Amoeboid cells can typically navigate these geometries without specific adhesions by expanding their leading edge through actin polymerization and coupling it to myosin-dependent contraction (40, 66, 67). Stratified epithelial tissues like the epidermis are an exception to this rule, as they lack a significant intercellular ECM component and are instead composed of densely packed and adherent cells (68, 69). Devices that impose planar confinement elicit stable bleb migration by amoeboid cells (18–21, 51), matching the mode of migration we observe in stratified epithelial tissues *in vivo*.

Stratified epithelial tissues like the epidermis may represent the clearest example of planar confinement a cell encounters *in vivo*. Indeed, our imaging of T cells in the epidermis of upright mounted zebrafish are highly reminiscent of T cells migrating in microchannels *in vitro* (51). While the basal and suprabasal keratinocyte layers are the confining substrates from the perspective of the T cell, previous work has demonstrated that basal keratinocytes are themselves under planar confinement between the basement membrane and the outer epidermal layers (70). Leukocytes may therefore experience planar confinement from the basement membrane to the most superficial cell layers of the epidermis. Whether T cells utilize this stable bleb mode of migration in other epithelial tissues and whether other amoeboid cells utilize stable blebs in epithelial tissues will help determine the universality of the tissue-specific migration programs we have described here. Further, it will be interesting to determine how inflammation-induced changes in keratinocyte mechanics (71) impacts confined leukocyte migration in the epidermis during infection and wound healing.

Finally, our characterization of zebrafish epidermal T cells strongly suggests they represent a DETC-like population, which is important for two reasons. First, it builds upon a growing body of work showing that intraepithelial T cells are evolutionarily ancient (72, 73). Whether these epidermal T cells regulate wound healing responses like they do in mice will be the subject of future investigation, but their presence in lampreys (72), teleost fish, and mammals (32) suggest an important and possibly evolutionarily conserved function. Second, larval T cell migration in zebrafish has been investigated as a way to study antigen surveillance (42), but our data indicates that migration at this developmental stage instead reflects tissue colonization. Because this tissue colonization occurs mostly *in utero* in mouse models, the external fertilization and imaging amenability of zebrafish may enable future discoveries on the formation, trafficking, and function of these developmentally early T cell populations.

## ACKNOWLEDGEMENTS

We would like to thank D. Bennin, A. Fister, G. Ramakrishnan, S. Shin, M. Mitchem, E. Townsend, N. Garcia and N. Mercado-Soto for their helpful discussion throughout the design and execution of this project. Figures throughout this paper were made in part with biorender.com. This study was funded by NIH grant R35 GM118027 (A.H.) and F32 GM146398 (T.F.R).

## Author contributions

T.F.R., J.S., and A. Huttenlocher designed research; T.F.R., J.S., and A. Horn performed research; Z.Z and J.R. designed and generated the tg(lck:lifeact-ruby) zebrafish; T.F.R. and J.S. analyzed data; A. Huttenlocher oversaw the project; and T.F.R. wrote the paper.

## MATERIALS AND METHODS

### Zebrafish husbandry and maintenance

All protocols using zebrafish in this study were approved by the University of Wisconsin-Madison Research Animals Resource Center (protocol M005405-A02). Adult AB strain fish and transgenic zebrafish lines including the *Casper* WT line (1) and previously published transgenic reporter lines *lck:*GFP (2), *cd4.1:*mCherry (3), *krt4*:lifeact-ruby (4) and *LyzC:*BFP (5) were used in this study. Double or triple transgenic reporters were generated by crossing individual lines and screening for fluorescence. For most experiments, these fish were then in-crossed to generate fish utilized for experiments. After breeding, fertilized embryos were placed into E3 media (5 mM NaCl, 0.17 mM KCl, 0.44 mM CaCl2, 0.33 mM MgSO4, 0.025 mM NaOH, and 0.0003% Methylene Blue) and maintained at 28.5°C in a petri dish in a laboratory incubator. *Lck:*GFP and *cd4-1:*mCherry larvae were screened at 6-7 dpf for fluorescence signal emanating from the paired thymus in the head. *LyzC:*BFP reporter fish were screened looking for fluorescent neutrophils in the caudal hematopoietic tissue anytime from 3-7 dpf. *Krt4*:lifeact-ruby reporter fish were screened by looking for fluorescent keratinocytes anytime from 3-7 dpf. At 7 dpf, screened larvae were transferred to a fish tank with still water where they were fed twice daily and maintained on a 14-h/10-h light/dark schedule. At 10 dpf, a drip of fresh water was applied to the tank, and at 21 DPF a gentle stream of water was applied. Fish were maintained at density of approximately 20 fish per 3-liter tank to maintain experiment-by-experiment growth consistency. Fish were removed from this environment as needed for experiments. When taking whole-animal confocal tiling images, fish were fasted overnight to reduce non-specific fluorescence coming from the intestinal tract.

### Cloning and DNA Injection for tg(*lck*:lifeact-ruby) zebrafish line generation

To construct Tol2:lck-Lifeact-Ruby primers (F: GCTAGCAAGATCTGCTCGAGCGAATTCACAATCTCATCATCATCATC R: GTGGCGAGATCCGGggtaccTTTACTAAGCATGAGAGAAAATGGT) were used to PCR Lifeact (Riedl, J. 2008.) and then in fusion cloning was performed to place Lifeact downstream of the lck promoter (p5E Lck was a gift from David Tobin; Addgene plasmid # 135201). 3 nL of solution containing 25 ng/μL of DNA and 35 ng/μL of Transposase mRNA was injected into the cytoplasm of one-cell stage unlabelled Casper or tg(*lck*:GFP) embryos. Fish were screened at 6-7 dpf for lifeact-ruby expression in the thymus, and fish with incomplete expression were sorted out for experiments. Fish with complete expression were used for establishing a stable tg(*lck*:lifeact-ruby) zebrafish line.

### Fish Handling and Scale Isolation for Imaging

For live imaging experiments, zebrafish larvae were added to E3 media containing 0.08 mg/mL tricaine (MS222/ethyl 3-aminobenzoate; Sigma-Aldrich). Once non-motile, warm 2% low melting point agarose (Thermofisher A-204-25) was mixed into 0.08 mg/mL tricaine E3 and anesthetized zebrafish were gently added to the bottom of a 50 mm glass bottom chamber (MatTek). Zebrafish were visually monitored as the agarose gelled and an eyelash brush was used to keep the fish flat during this process. In some experiments (Fig. 6), fish were deliberately maintained in the upright position during this process with an eyelash brush. When handling juvenile or adult zebrafish (Fig. 1), anesthetized fish were moved to 1-well (Nunc Lab-Tek II 155360) glass-bottom imaging chambers with a transfer pipet (Fisher 13-711-7M) cut halfway to the base to increase the hole size. For tiling experiments, warm 2% agarose was then added to the chamber and allowed to polymerize prior to imaging. For live adult imaging (Fig. 1), drift was then limited by adding soaked sponge cloth to the chamber around the fish, and an additional soaked sponge cloth was draped over the fish head and upward-facing gills to maintain viability. In all cases, the fish were imaged for ≤20 min. Following imaging, the fish were killed by keeping them in 0.16 mg/mL tricaine for 20 min. For nonterminal experiments, the fish were quickly transferred to a tank of fresh RO water to promote recovery following imaging experiment. For scale explant experiments, anesthetized adult fish were transferred to a sponge cloth under a light dissecting microscope (Nikon SMZ-745) and fine point tweezers (Dumont) were used to pluck scales, which were transferred to a dish with imaging media (Leibovitz’s L-15 Medium without phenol red supplemented with 2% fetal bovine serum and 1X Gibco Penicillin-Streptomycin). Grouped scales (intact) were gently transferred a 4-well (Ibidi 80427) glass bottom chamber. For the wounded group, scales were gently teased apart into individual scales and forceps were firmly pressed through the region of the scale covered with epidermis prior to transfer. For all inhibitor experiments, scales were incubated with the indicated concentration of para-amino-blebbistatin (Cayman 22699), Y-27632 (Fisher 125410), or an equivalent volume of dimethylsulfoxide (DMSO; Sigma-Aldrich D2650) for at least 20 minutes prior to imaging. To visualize scales, we immersed the euthanized fish into 0.04% solution of Alizarin red powder (Sigma Aldrich 130-22-3) dissolved in RO water for 20 min. The fish were then washed for three times for 5 min in fresh RO water with gentle rocking prior to imaging. For puncture wounding (Fig. 6), A 30G½ needle (Becton Dickinson PrecisionGlide) was used to puncture the caudal peduncle on the dorsal side of the animal. Pressure was applied until the needle came out the other side. The wounded fish were recovered in a tank of fresh system water until movement was observed (0 to 3 min) and then maintained for 2 hours until imaging.

### Microscopy and Image Preparation

For low magnification images of the intact and wounded scales, scale explants were imaged on a Zeiss Zoomscope (EMS3/SyCoP3; 1× Plan-NeoFluar Z objective; Zeiss) with an Axiocam Mrm charge-coupled device camera using ZenPro 2012 software (Zeiss). For all other fish imaging, we used a spinning disc confocal microscope (CSU-X, Yokogawa, Sugar Land, TX) with a confocal scanhead on a Zeiss Observer Z.1 inverted microscope, either an EC Plan-Neofluar 10×/0.30 or EX Plan-Neofluar 40×/0.75 object, a Photometrics Evolve EMCCD camera, and Zen software (Zeiss). To stitch tiled images in ZenPro 2012 software (Zeiss), we first performed a max intensity z-projection and then stitched with a 10% maximal shift. All images were prepared for publication with ImageJ (v. 2.1.0/1.53c).

### Cell Tracking and Analysis

Cell migration movies to be tracked were imported into ImageJ (v. 2.1.0/1.53c), and individual cells were tracked using the Manual Tracking or TrackMate plugin. For calculating T cell velocity over development, 10 randomly selected cells per larvae were manually tracked from each movie. For calculating the lifeact-ruby mean gray value (Fig. S2), every analyzable cell was outlined using the wand tool and measured in ImageJ. For calculating the leading edge to trailing edge max intensity ratio (Fig. 3/Fig. 4), the front and rear 1/3^rd^ of the cell were manually outlined, and the lifeact-ruby integrated density was measured for each site. The local background for each cell was measured and subtracted from the leading and trailing edge values prior to calculating the ratio. For calculating the aspect ratio (Fig. 3), cells from each site were outlined with the wand tool and measured in ImageJ. For correlating actin spatial distribution with cell velocity (Fig. 4), instantaneous LE/TE ratio was correlated with velocity in the 5-15 surrounding frames for 45 total cells (3 independent scale explant experiments obtaining 3 movies per experiment and tracking 5 cells per movie). Measurements of cell-front and actin flow velocity (Fig. 5) were made from kymographs (x-axis denoting distance; y-axis representing time) of cell migration (lck:lifeact-ruby) as the inverse tangent of the angle between lifeact signal and the y-axis. Measurements were made over 3 independent experiments, each tracking 5 independent cells with mean velocities determined from 1-3 measurements per cell. Average T-cell velocities under treatment with blebbistattin, Y27632 or DMSO (Fig. 5) were quantified using the automated TrackMate plugin in FIJI/ImageJ over 3 independent experiments with 2 independent movies per experiment per condition.

### Statistical Analysis

Statistical tests were all performed using GraphPad Prism (version 10.2.3), with specific tests and p-values indicated in each figure legend.

