## Supplemental Figures for "Live imaging in zebrafish reveals tissue-specific strategies for amoeboid migration"

**Figure S1. T cells enter the tail fin primarily through the caudal vein.**

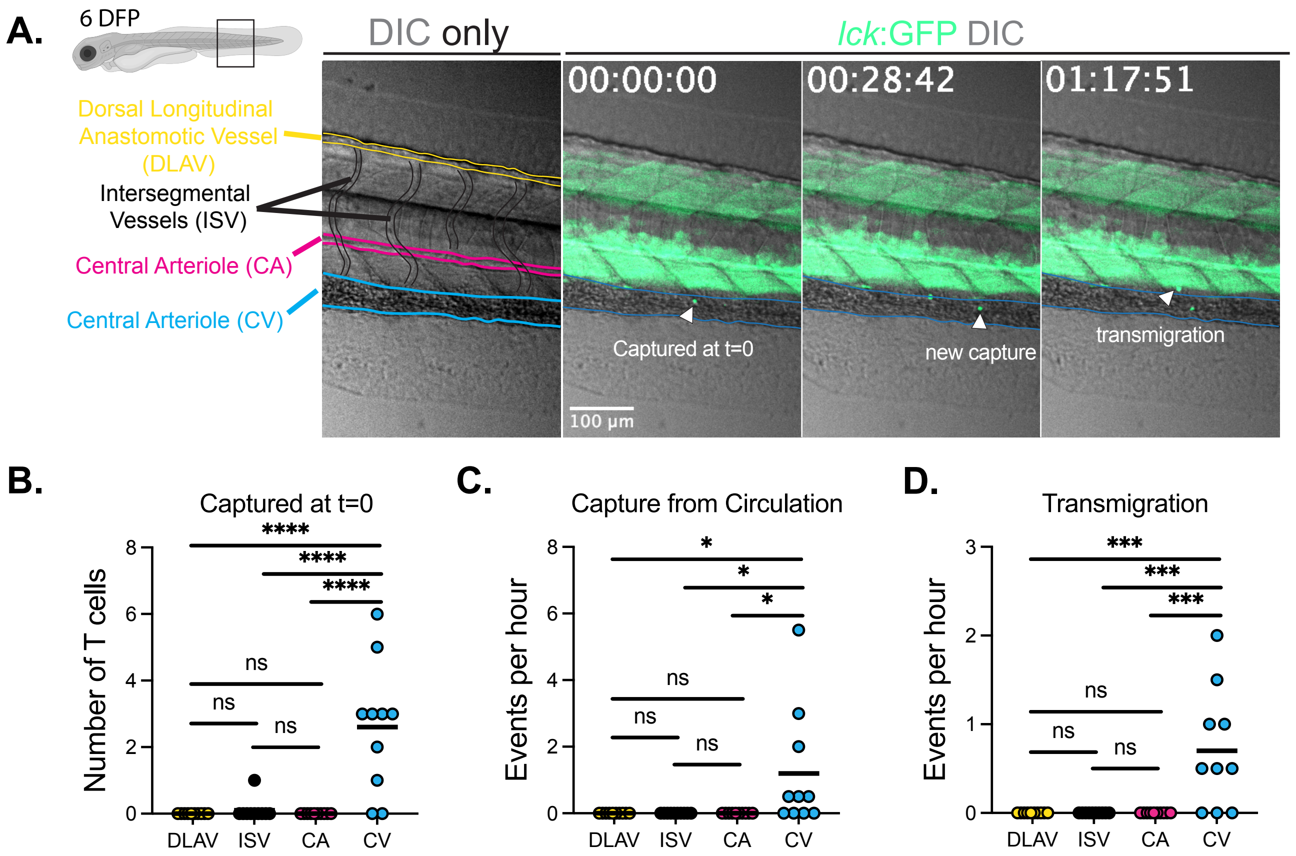


**Supplemental Figure 1. T cells enter the tail fin primarily through the caudal vein.**
**A.** Timelapse imaging of a 6 dpf tg(lck:GFP) zebrafish with a labelled image showing the different blood vessels in the caudal peduncle. Note that vessels are more apparent in movies where blood flow is apparent. Examples of a T cell being capture at the start of the movie (t=0), being captured from circulation, and transmigrating into the tissue are noted. Scale bar = 100 microns. **B-D.** Quantification of which blood vessels T cells are initially captured on (B), which blood vessels they are captured onto over the course of 1-2 hour movies (C), and which vessel they transmigrate through to enter the tissue (D). In each graph, means are compared with an ordinary one-way ANOVA. P values were denoted on graphs as follows: n.s., $P > 0.05$; *, $P < 0.05$; **, $P < 0.01$; ***, $P < 0.001$; ****, $P < 0.0001$.

**Figure S2. lifeact intensity is lower in peripheral tissues relative to the thymus**

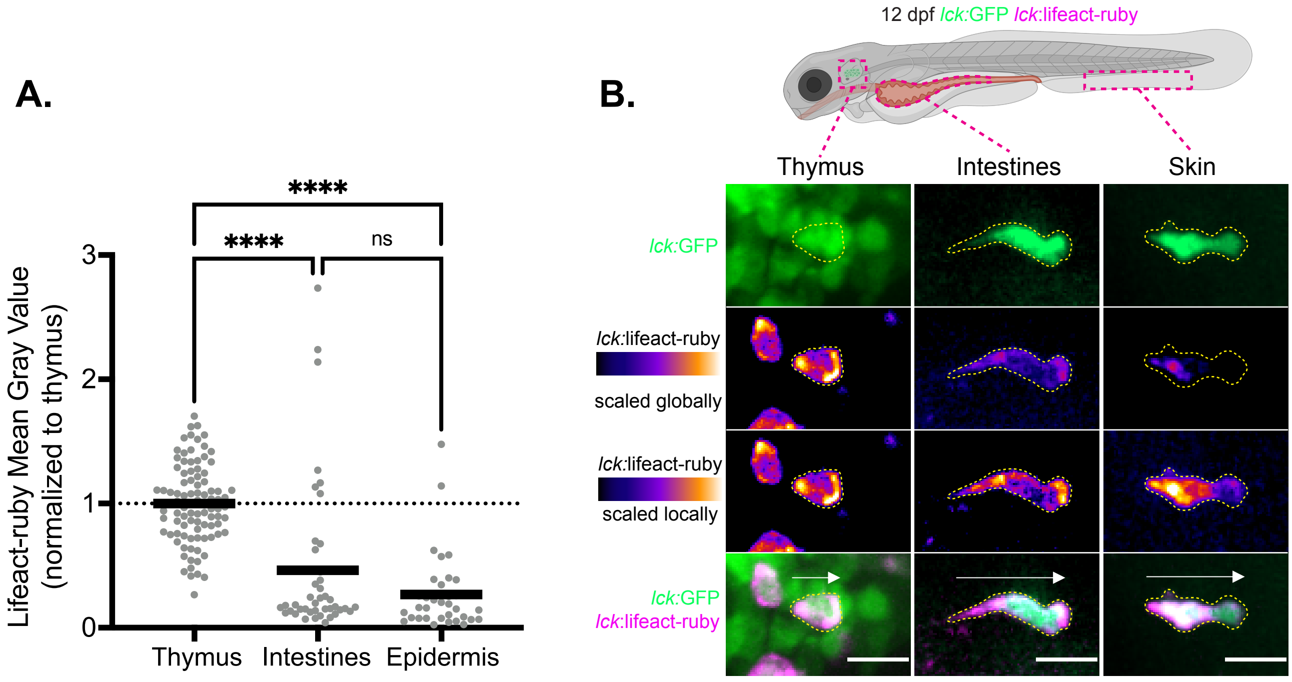


**Supplemental Figure 2. Lifeact intensity in T cells across the body**
**A.** Quantification of the mean gray value of lifeact-ruby in T cells from the thymus, intestines, and epidermis normalized to the average thymus value. Each dot represents a cell pooled from 3 independent experiments. The means were compared with an ordinary one-way ANOVA. ****$p<0.0001$. **B.** Representative lifeact-expressing T cells from the thymus, intestines, and epidermis all scaled the same (globally) or all scaled individually (locally). In the merged image, the direction of migration and cell outline are noted. Scale bars = 10 microns.

**Figure S3. Intact and forceps-wounded scale explants**

**A.**

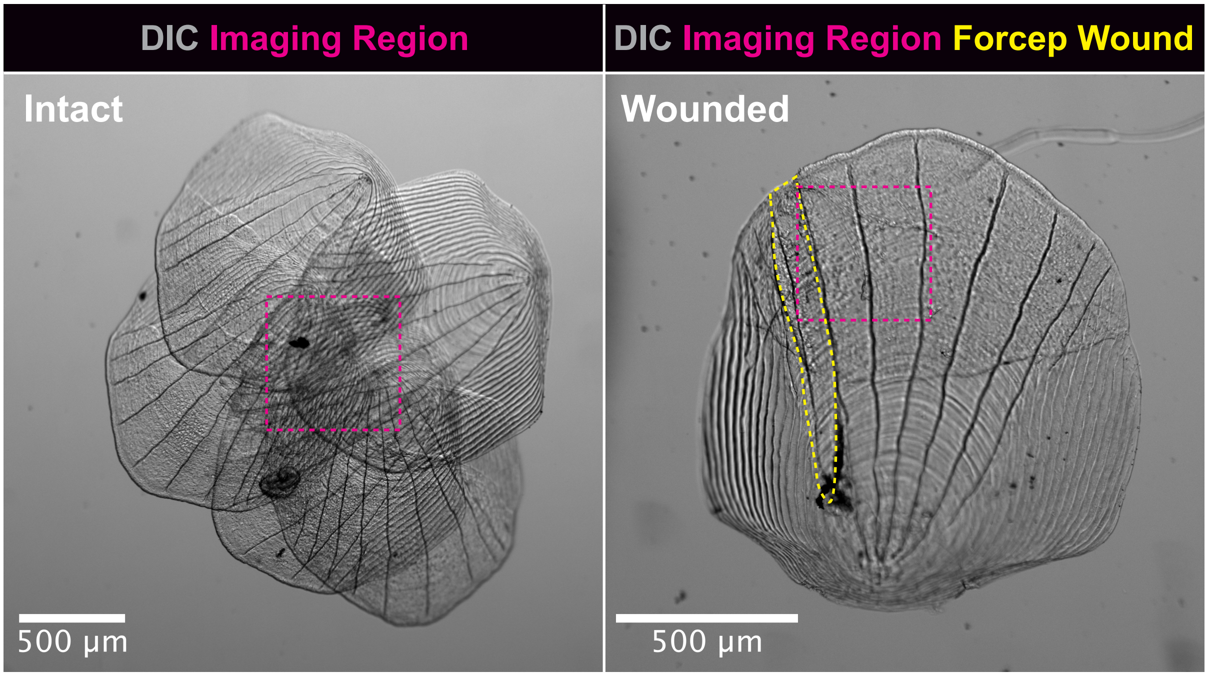


**Supplemental Figure 3. Intact and forceps-wounded scale explants.**
**A.** Low magnification DIC imaging of intact scale clusters and forceps-wounded scales with the imaging region highlighted in magenta and the forceps wound highlighted in yellow. Representative of 2 individual experiments. Scale bars = 500 microns.

**Figure S4. The actin-poor leading edges in epidermal T cells are stable**

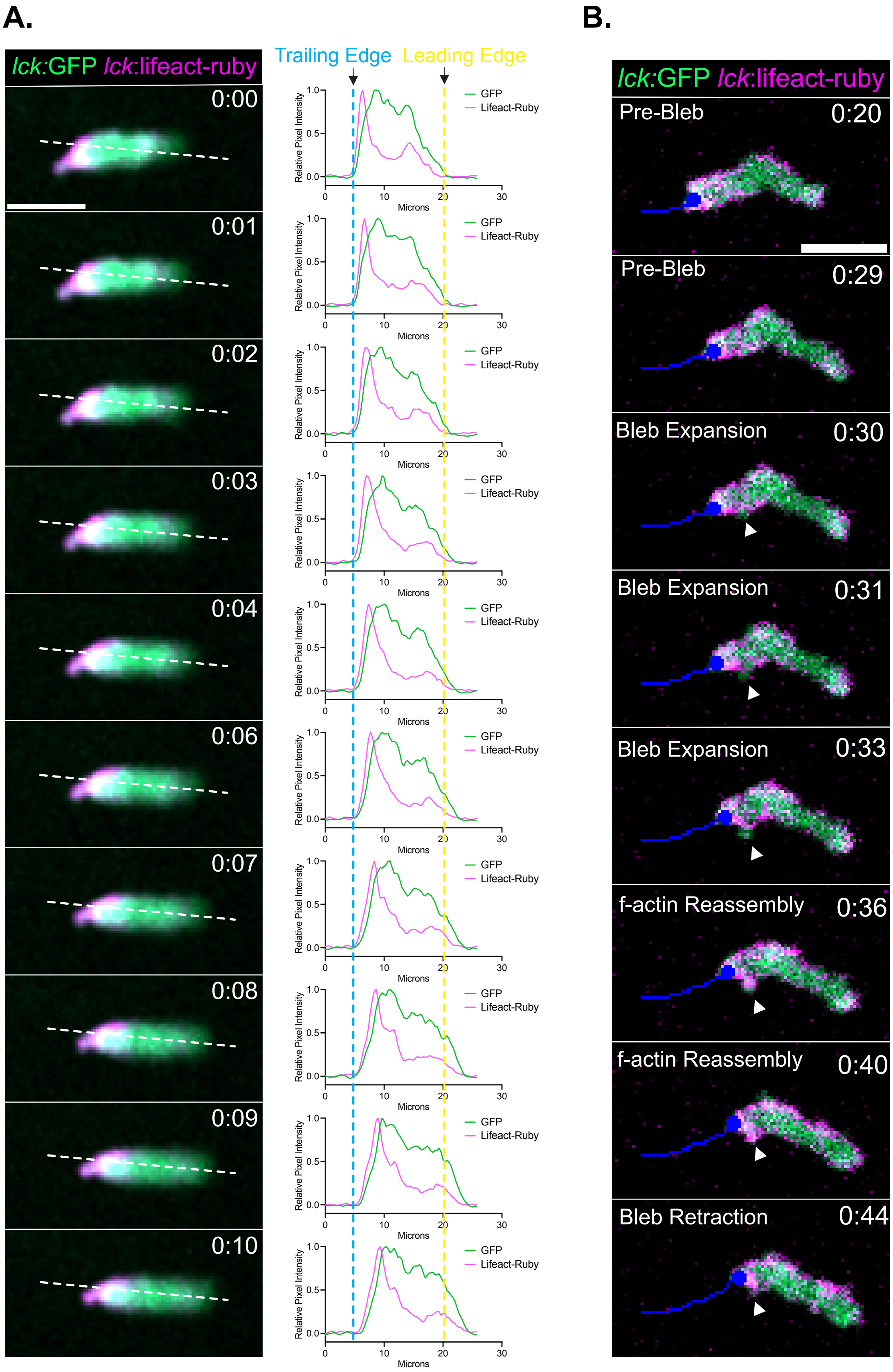

**Supplemental Figure 4. The actin-poor leading edge in epidermal T cells is stable.**

**A.** Rapid acquisition timelapse images of epidermal T cells from 12 dpf tg(lck:GFP; lck:lifeact-ruby) larvae and associated line profile analysis showing the steady progressive movement of the f-actin-poor cytoplasmic leading edge. Time in mm:ss. **B.** Rapid acquisition timelapse images of rare intermittent bleb expansion and retraction in an epidermal T cell noted by arrowhead. Time in mm:ss. Scale bars = 10 microns.
